# A robust relationship between robustness and evolvability

**DOI:** 10.1101/2024.07.08.602504

**Authors:** Nate B Hardy

**Affiliations:** 301 Funchess Hall, Department of Entomology and Plant Pathology, Auburn University, Auburn, Alabama, 36849

**Keywords:** gene-by-gene interactions, epistasis, adaptive potential, canalization

## Abstract

Here, I consolidate and extend simple models of how genetic robustness affects the evolvability of phenotypes with discrete states. I share three main insights. (1) What we already know about the effects of mutational robustness on evolvability can be readily transferred to environmental robustness. (2) Even without a hard genotype-level trade-off between robustness and evolvability, the optimal level of phenotypic robustness depends on the rate of environmental change. But counter-intuitively, when adaptive landscapes are complex, an increase in environmental stability can increase the frequency of environmentally-robust but mutationally-sensitive genotypes. (3) Even with a fixed probability that a mutation is neutral, populations can evolve along the spectrum of robustness and evolvability by evolving the genotype-determined neighborhood of mutationally-accessible phenotypes. Indeed, because it allows for the evolution of increased evolvability without a concomitant increase in genetic load, selection should favor changes in the phenotypic neighborhood over changes in mutational sensitivity.

**Teaser Text:** Models of the evolution of discrete phenotypes show that their evolvability is highest when environments fluctuate at intermediate rates, and with intermediate levels of mutational robustness. By merging previous models and relaxing previous assumptions, I show that previous insights are robust. I also show that the same models tell us very different things about the evolvability of discrete traits, namely, that what applies to robustness against mutation is readily transferable to robustness against environmental perturbation, and that selection for evolvability will preferentially work on the neighborhood of mutationally-accessible phenotypes rather than the probability of non-neutral mutation.

## 1. Introduction

Phenotypes tend to be robust against much genetic and environmental variation (1,2). If the targets of selection are stable, such robustness can be adaptive, as it promotes homeostasis and decreases genetic load (3–6). On the other hand, robustness reduces phenotypic variation (7) so, if the adaptive landscape varies, selection may promote genotypes that are more sensitive to mutational or environmental variation, and are thus more evolvable (8–10).

Genetic and environment variation map to phenotypic variation via developmental systems that can be complex, and fittingly, many studies of the relationship between robustness and evolvability have used complex models of such mappings (11,12). Examples include empirically-based models of RNA or protein folding (7,10,13), or more theoretical models of self-assembling systems (14,15) or gene regulatory interactions (16–19). Although apropos, the interpretation of such models is fraught due to the fact that robustness is not the only thing affected by structural variation in a genotype-phenotype map. In particular, such variation can also affect how a developmental system (1) accumulates cryptic genetic diversity (20), (2) phenotypically responds to non-neutral mutation (21,22), (3) transitions between ordered and chaotic dynamics (19), and (4) “learns” from past selective environments (22–24). So, with complex genotype-phenotype models, when a change in evolvability is observed, it can be difficult to say that a change in robustness *per se* is the cause.

An alternative approach is to abstract away the details of a developmental system, and model robustness simply as the probability that a change in the genetic or external environment will affect a change in phenotype (25,26). Such models have shown, somewhat counter-intuitively, that evolvability can be facilitated by intermediate levels of robustness (25), and promoted by intermediate rates of environmental change (26). But the extent to which these insights depend on specific model assumptions is unclear and therefore so are the boundaries of the theory’s domain of applicability. Our goal here is to relax those assumptions, expose heretofore unrecognized connections between models, and get a better sense for when phenotypic robustness is good for the evolvability of discrete traits. Let us start with a quick look at the work that provides the foundation for what follows.

## 2. Simple models of the robustness of discrete phenotypes

### 2.a. Genotype robustness

At the genotype level, robustness and evolvability trade off; robustness decreases phenotypic variance (10). Meyers et al. (26) elegantly demonstrate that given genotype-level trade-offs between mutational robustness, environmental robustness, and evolvability (10), the optimal level of genetic robustness is a function of the rate of environmental change. They ask us to imagine a pentagonal, bidirectional, mutational network, where each vertex is a genotype, and each edge is a mutation (Fig. 1a.). A population of genotypes that can mutate along the edges of this network is exposed to fluctuating selection. Each genotype maps to one of three possible phenotypes: (1) phenotype *A* is optimal when the environment is in state *a*, (2) phenotype *B* is optimal when the environment is in state *b*, and (3) phenotype *V* has intermediate fitness in both environmental states (Fig. 1a). To clarify how this model relates to the next we will consider, let us define a couple new terms. First, let *q_i_* denote the probability that a mutation of genoptype *g_i_*has no effect on the phenotype. In Fig. 1a, *q_i_* varies from one for genotype *g_1_,* to zero for *g_3_* and *g_4_*. Second, let *r_i_* stand for the environmental robustness of genotype *g_i_*, that is, the probability that a change in the environment has no effect on fitness. In Fig. 1a, it is one for *g_3_* and less than one for the rest. Mutational and environmental robustness are both genotype-specific.

**Figure 1.**
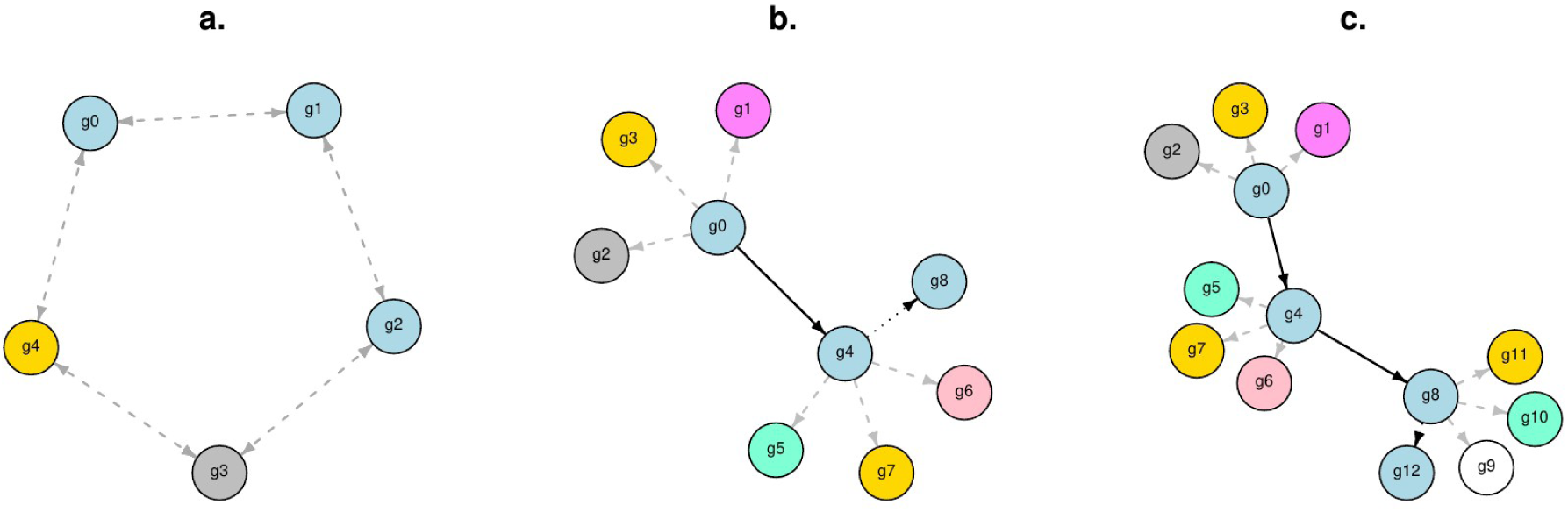
Mutational networks with redundant mappings of genotypes to phenotypes. (**a**) The network of Meyers et al. (26). Five genotypes are possible, each of which maps to one of three phenotypes: A (light blue) is optimal in state a environments; B (gold) is optimal in state b environments; V (gray) has intermediate fitness across environmental states a and b. Back-mutation is possible and the graph forms a cycle. Robustness varies across genotypes, but the robustness and phenotypic neighborhood of each genotype does not change over time. (**b, c**) Example of the mutational network of Draghi et al. (25) at two time points. Different vertex colors denote different phenotypes, with the phenotype p_0_ (light blue) being currently optimal. Potential phenotype-changing mutations are shows with dashed gray edges; the set of vertices connected by such edges to a genotype constitute that genotype’s phenotypic neighborhood, **k_i_**. Potential neutral mutations are shown as dotted black edges, and realized neutral mutations are shows as solid black edges. Back-mutation is forbidden, thus, the graph is divergent.

Meyers et al. (26) show that given this mutational network, the expected frequency of each genotype depends on a parameter *ƛ,* the number of generations between a change in the selective environment (Fig. S1): (i) When the environment changes infrequently (*ƛ* > 1000) the most mutationally robust genotype, *g_1_*, occurs at the highest frequency. (ii) When the environment changes rapidly (*ƛ* < 10), the most important thing is the environmental robustness of a genotype, *r_i_*; the generalist genotype *g_3_* occurs at the highest frequency (8). (iii) When the environment changes neither too fast nor too slow (10 < *ƛ* < 1000), the population evolves to maximize evolvability; the highest frequency genotype is *g_0_*, which is the genotype that maps to phenotype *A* that has the lowest combination of mutational and environmental robustness. In sum, selection works not just on the observable phenotype {*A*, *B*, *V*} but also the robustness and evolvability of those phenotypes.

### 2.b. Phenotype robustness

At the phenotype level, robustness and evolvability need not trade off, since robustness can increase cryptic genetic diversity (29,30,31,32), an elegant demonstration of which is given by Draghi et al. (25). Like Meyers et al. (26), they consider an evolving population of genotypes the relations of which can be visualized as a mutational network (Fig 1b,c). But in this case, instead of specifying a fixed, cyclical network, they specify a mutational process, and allow a divergent network of mutations to emerge organically. They let each genotype *g_i_* map to a phenotype *p_i_*, and have a *K*-dimension set **k_i_** of alternative phenotypes that can be reached by a single mutation. Thus, **k_i_** is what has been referred to as the phenotypic 1-neighborhood of a genotype (10). (Note that in the Meyers et al. (26) model, each genotype also has a phenotypic neighborhood, but there is no requirement that adjacent genotypes have alternative phenotypes, and thus *K* varies across genotypes.) Phenotypes for *p_i_* and **k_i_** are chosen randomly from a set of possible phenotypes **P** of size *P*. Before time *t_x_*, a population of genotypes is exposed to strong stabilizing selection about an initial phenotype *p_0_*; all alternative phenotypes confer a fitness of zero.

Mutations at rate *μ* are neutral with probability *q*; thus, as in the model of Meyers et al. (26), *q* affects mutational robustness, but in this case *q* does not vary across genotypes. At rate *μ*(1-*q*), mutation causes a change in the phenotype, with the new phenotype, *p_i_’,* selected from the phenotypic neighborhood **k_i_**. Whenever a mutation occurs, regardless of whether or not it is neutral, a new phenotypic neighborhood, **k_i_**, is randomly sampled from the phenotypic space **P**. So, during this initial phase of evolution, while strong stabilizing selection prevents a population from diversifying phenotypically, the population can grow a network of phenotypically redundant genotypes, each of which has its own phenotypic neighborhood (Fig. 1b,c).

At time *t_x_*, a new optimal phenotype, *p_x_*, is chosen from **P**. Draghi et al. (25) examine how, *t_a_*, the time it takes for the population to evolve *p_x_* depends on *q*, *K*, and *P*. They show that if *K* = *P –* that is, every genotype can with one mutation reach every possible phenotype – increasing mutational robustness *q* monotonically increases the expected time for the population to evolve *p_x_*(Fig. 2). As in Meyers et al. (26), robustness and evolvability are antithetical. But if *K* < *P*, mutational robustness can increase the diversity of phenotypic neighborhoods, and the relationship between *q* and the expected time for the evolution of *p_x_* is non-monotonic; evolvability is highest at intermediate values for *q* (Fig. 2).

**Figure 2.**
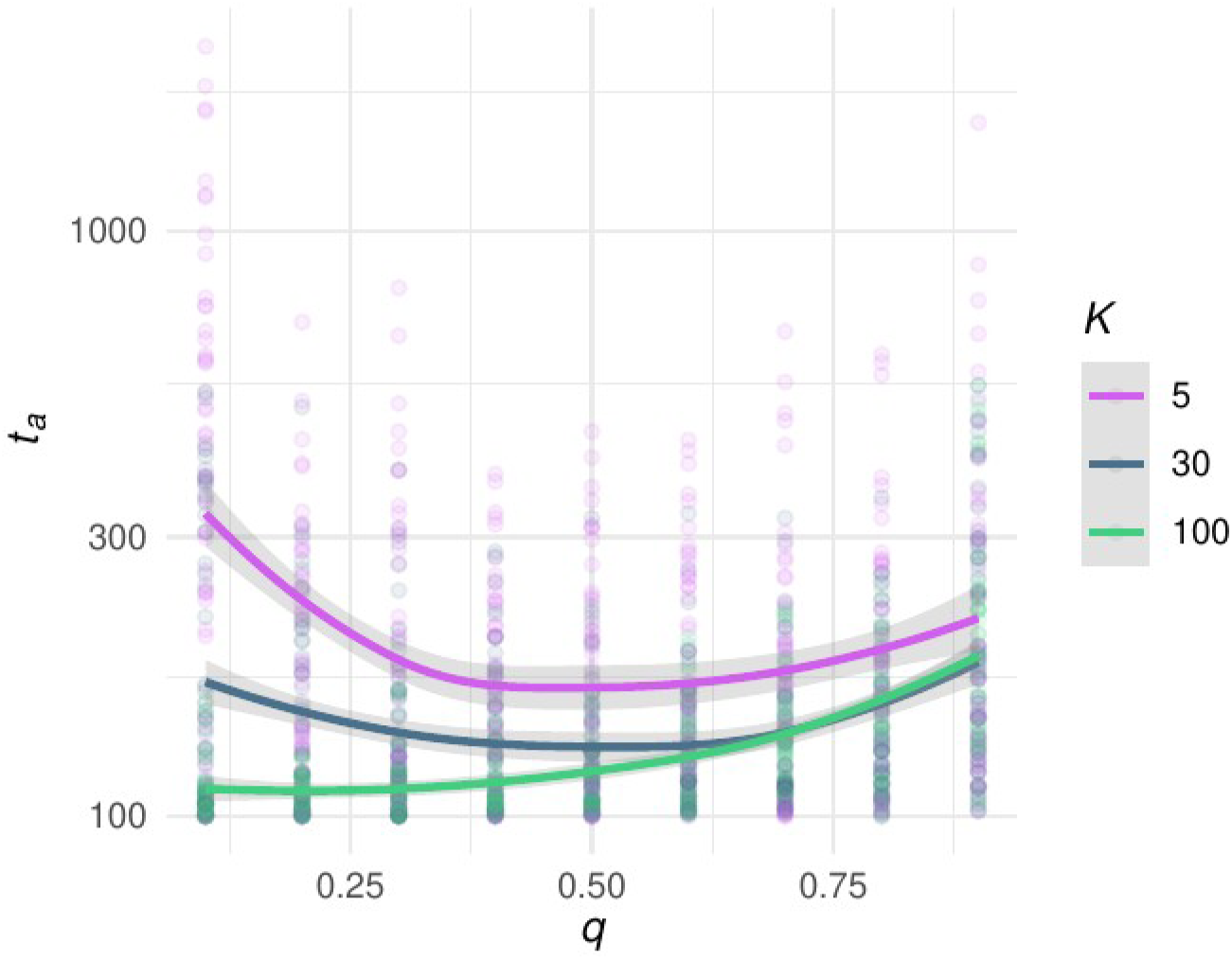
The effect of mutational robustness, q, on evolvability, t_a_^-1^, depends on the difference between the number of possible phenotypes, K, and the size of the phenotypic neighborhood P. This reproduces the results of Draghi et al. (25) but with an individual-based simulation model (Supplementary File S2), with 50 replications for each combination of parameter values, P = 100, K = {5,30,100}, and q = (0.1, 0.2, …, 1.0). The y-axis is on a log scale. Curves are loess regressions, with a span of 1.0; shaded areas give 95% confidence intervals. Interpretation*: Unless all phenotypes are accessible by one mutation, evolvability is maximized at intermediate levels of mutational robustness*.

## 3. The evolution of environmental robustness

Although Draghi et al. (25) and Meyers et al. (26) focus on mutational robustness, their models are readily transferrable to questions about environmental robustness.

Imagine that, as in the Draghi et al. (25) model, there is a world with *P* possible phenotypes. But instead of a population of genotypes that explores this world via mutation, suppose there is a meta-population of genetically-invariant demes that explores it via migration. Phenotypes emerge from genotype-by-environment interactions, and environments vary such that each deme has a *K*-dimensional phenotypic neighborhood, **k_i_**, accessible via migration. For a depiction of this model, Figure 1.b. should suffice; one need only reinterpret the nodes as demes, and the edges as migration events. Let *q_m_* represent the probability that the environment of a new deme induces no change of phenotype; in other words, *q_m_* determines the migrational robustness of the genotype to environmental variation. Since we assume the population is genetically monomorphic and have replaced mutation with migration, we can not assess the relationship between robustness and evolvability *per se,* at least not if we follow convention and think about evolution only in terms of changes in genotype frequencies. But we can assess the relationship between robustness and plasticity, which is analogous. In the beginning, only phenotype *p_0_*is adaptive and only demes that express *p_0_* survive selection. But migration forms a neutral network of demes, and the diversity of migrationally-accessible phenotypic neighborhoods grows. If at some point a new phenotype, *p_x_*, becomes optimal, as long as *K* < *P*, we know that the relationship between *q_m_* and rate at which the population evolves via migration to express *p_x_* will be non-monotonic, with the fastest rates occurring at intermediate values of *q_m_*. This is because the logic of Draghi et al. (25) model of mutational robustness has not changed, only our interpretation of some of its variables and parameters. Neutral mutation can boost evolvability by changing the structure of a mutation network. Neutral migration can boost plasticity by changing the structure of a population in a heterogeneous environment.

To go beyond plasticity, and connect environmental robustness to evolvability, we need to allow for mutation as well as migration (Fig. S3). Above, we let phenotypes vary depending on the interaction between one genotype and many local environments. Suppose instead that each individual can have one of two genotypes, either *g_0_*or *g_1_*. Initially, the population is monomorphic for *g_0_*but mutation between genotypes occurs at rate *μ* per individual per generation. In *e_0_* environments, genetic variation is cryptic; both genotypes express the optimal phenotype *p_0_*. Initially, migration to any non-*e_0_* environment induces strongly deleterious changes in phenotype. Then at time *t_x_*, a new phenotype *p_x_* – which is expressed only by *g_1_* genotypes in *e_x_* environments – becomes adaptive in *e_x_* environments. Now we can ask the question that ties environmental robustness to evolvability: How does the time it takes a population to evolve phenotype *p_x_* depend on environmental robustness, *q_m_*? Analysis of this model (with codes given in Supplementary Document S5) shows that evolvability is highest at intermediate values for *q_m_* (Figure S4). Robustness is robustness; mutational and migrational robustness have similar effects on evolvability.

In a similar fashion, the model of Meyers et al. (26) can be transferred to the evolution of environmental robustness and evolvability; we need only change our interpretation of the edges and vertices in Fig. 1a. Instead of five genotypes, we have one phenotypically-plastic genotype, five spatial locations, and three different environment types, such that the genotype can expresses three different phenotypes depending on its environment. Instead of genotypes connected by mutation, we have demes connected by migration. If we apply the same kind of fluctuating selection – with the phenotype expressed in demes *g_0_*, *g_1_*, and *g_3_* being optimal in temporal state *a*, that expressed by deme *g_4_* being optimal in state *b*, and that in *g_3_* with intermediate fitness in both states – then the proportion of a meta-population in each deme will be a function of the rate of environmental change. When environmental change is slow, individuals will occur at the highest frequency in deme *g_1_*, since that minimizes migration load (29). When the world turns over rapidly, deme *g_3_* offers a safe harbor of stability. And at intermediate rates of change, the highest fitness can be realized by hopping back and forth between demes *g_0_*and *g_4_*.

## 4. Consolidating models

In Sections 2 and 3, I pointed out fundamental similarities between the models of Meyers et al. (2005) and Draghi et al. (2010). In this section, I merge those models and relax some their main assumptions. Doing so shows that their main inferences are robust. It also reveals some more nuanced dynamics. I take an individual-based approach, with models developed using the SLiM framework (30)

### 4.a Divergent mutational networks of genotypes varying in mutational and environmental robustness

#### 4.a.i. Model description

Imagine a population of haploid individuals in an environment with a carrying capacity of *N*=500, and for which there is a *P-*dimensional set **P** of possible phenotypes. Each individual has three genetically-determined traits: (1) a phenotype *p_i_*, (2) a phenotypic neighborhood **k_i_**of size *K*, and (3) a mutational robustness *q_i_*. The life cycle follows the Wright-Fisher model, with selection on reproductive success, clonal reproduction with mutation, and non-overlapping generations. At the start of each simulation, the population is monomorphic, with *q_i_*= 0.1, and an initial phenotype *p_0_*and phenotypic neighborhood **k_0_** selected randomly from **P**. Mutation happens at rate *μ*=2e-3 per individual per generation. Each mutation is neutral with probability *q_i_*. Whenever a neutral mutation occurs, a new phenotypic neighborhood **k_i_** is chosen from **P**. When a non-neutral mutation occurs, there is an equal chance that it affects either mutational robustness *q_i_*, or the phenotype *p_i_*. If the former, a new value for *q_i_*is chosen from a random uniform distribution. If the latter, a new phenotype *p_i_* and phenotypic neighborhood **k_i_** are randomly sampled from **P**.

This population is subjected to fluctuating selection at rate *ƛ^-1^*. When the environment is in state *a*, phenotypes in *L*-dimensional set **O_a_** are optimal, and when the environment is in state *b*, a non-overlapping *L*-dimensional set **O_b_** of phenotypes are optimal. Phenotypes in *L*-dimensional set **V** have intermediate fitness in both environmental states. So,

*w_a_*(*p_i_*) = *β_0_*+*s* for *p_i_* ∈**O_a_**; *β_0_* for *p_i_* ∈ **O_b_**; *β_0_*+*sh* for *p_i_* ∈ **V**, and

*w_b_*(*p_i_*) = *β_0_* for *p_i_* ∈**O_a_**; *β_0_*+s for *p_i_* ∈ **O_b_**; *β_0_*+*sh* for *p_i_* ∈ **V**

where, similar to Meyers et al. (26), *w_j_* gives the fitness of phenotype *p_i_* in environmental state *j*, *β_0_* = 0.6 is the base reproductive fitness, *s* = 0.4 is the fitness advantage of expressing an appropriate specialist phenotype, and *h* = 0.5 determines the intermediacy of the generalist phenotype.

We simulate evolution by repeating this life cycle for 20K generations, keeping track of the following model variables: (1) *w*, the mean population fitness, (2) *q_i_*, the mean population mutational robustness, (3) *D*, the mean Simpson’s Diversity of the population’s phenotypic neighborhoods, and (4) *f_V_*, the mean frequency of individuals expressing a generalist phenotype, that is, *p_i_* ∈ **V**. Fifty replicated model simulations were run for every combination of model parameters (Table 1). Model codes are provided as Supplementary Document S3.

**Table 1.** Divergent network model parameters and variables.

11 Genetic Robustness and Evolvability
| Parameter | Description | Values |
| --- | --- | --- |
| $\mu$ | Mutation rate | 2e-3 |
| $N$ | Population size | 500 |
| $P$ | Number of possible phenotypes, **P** | {6, 100} |
| $K$ | Size of phenotypic neighborhood, **k<sub>i</sub>** | {1, 3, 6} \| $P=6$<br>{6, 30, 100} \| $P=100$ |
| $L$ | Number of phenotypes in each set **O<sub>a</sub>**, **O<sub>b</sub>**, and **V** | 1 \| $P=6$<br>$K/3$ \| $P=100$ |
| $\lambda$ | Number of generations between change in environment | {1e+1... 1e+4} |
| $\beta_0$ | Viability of specialist genotype in wrong environment | 0.6 |
| $s$ | Viability advantage of specialist in right environment | 0.4 |
| $h$ | Viability intermediacy of generalist phenotype | 0.5 |
| Variable | Description | Range |
| $w$ | Population mean fitness | $0 < w < 1$ |
| $q_i$ | Population mean mutational robustness | $0 < q_i < 1$ |
| $f_v$ | Frequency of generalist phenotypes ($p_i \in \mathbf{V}$) | $0 < f_v < 1$ |
| $D$ | Simpson's Diversity of phenotypic neighborhoods, **k<sub>i</sub>** | $0 < D < 1$ |

#### 4.a.ii Model Dynamics

First consider what happens when the set of possible phenotypes is small (*P*=6) and only one phenotype is optimal in each environmental state (*L*=1) (Fig. 3). Regardless of phenotypic neighborhood size, *K*, when selection fluctuates rapidly (*ƛ* = 10) mean population fitness is ∼ 0.8 (Fig. 3a), which is the viability of the generalist phenotype, and indeed, the generalist phenotype tends to occur at frequencies near one (Fig. 3c). As the rate of fluctuating selection decreases, mean population fitness rises, and the frequency of the generalist phenotype falls, so that when *ƛ* =10,000, *w* is near one, and *f_V_* is near zero. (But note that there is a kink in the mapping of *ƛ* to *f_V_*, at ∼ *ƛ*=100; we will return to this below.) In contrast, the relationships between *ƛ* and the other model statistics, *q_i_* and *D*, are non-monotonic. Mutational robustness, *q_i_*, is lowest at intermediate values for *ƛ*, with the bend of the curve being more pronounced with larger phenotypic neighborhoods (Fig. 3b). We see a similar U-shaped mapping of *ƛ* to the mean diversity of phenotypic neighborhoods, *D*, although when *K*=*P*, *D* equals one by definition, and the extent to which *D* drops at intermediate values of *ƛ* increases with decreasing *K*. If populations are more mutationally sensitive and hence more evolvable, repeated bouts of positive selection will tend to sweep away much of the standing diversity in phenotypic neighborhoods (31). This effect is especially strong when *K*=1, in which case it takes longer for *D* to rebound from a sweep. So, when *K* ∈{3, 6} and *P*=6, increased mutational sensitivity (1-*q_i_*) clearly brings about increased evolvability. On the other hand, when *K*=1, this relationship weakens, as a relatively small dip in mutational robustness, *q_i_,* comes with a relatively large dip in the diversity of phenotypic neighborhoods, *D;* though more mutations are non-neutral, they access a smaller part of the phenotypic space, *P*.

**Figure 3.**
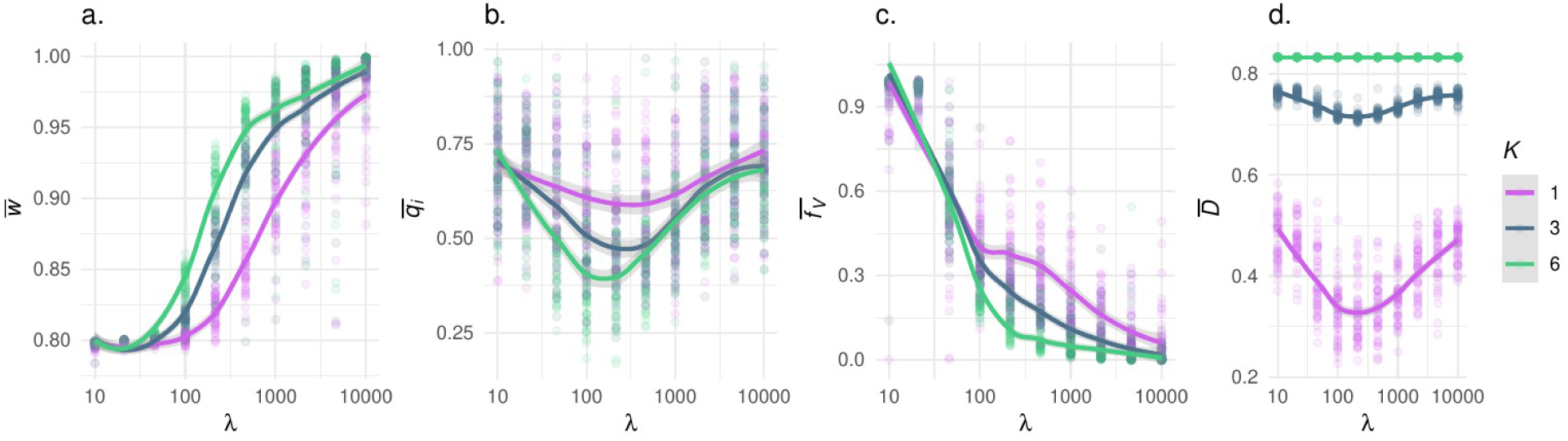
Summary of models where K is fixed and q_i_ is freely evolvable. Panels show how the number of generations between environmental fluctuations, ƛ, affects (**a**) population mean fitness, w-bar, (**b**) mean mutational robustness, q_i_-bar; (**c**) mean frequency of generalist genotypes f_V_-bar, (**d**) mean population diversity of phenotypic neighborhoods, D-bar. Each point shows the result for one simulation. Lines show loess regressions. Point and line colors denote different values for phenotypic neighborhood size, K. Interpretation*: As environments become increasingly stable, there is a succession of mutationally robust generalists, evolvable specialists, and mutationally robust specialists*.

When the set of possible phenotypes is comparatively large (*P*=100) and more than one phenotype is optimal in each environmental state (*L*=*K*/3), what we find is similar to when *P*=6 and *L*=1 (Fig. 4), but when *K=*6, the predominance of specialist genotypes in more stable environments is much less pronounced, and the kink in the mapping of *ƛ* to *f_V_* is yet more striking. (Fig. 4a). In fact, polynomial regression analysis recovers support for a cubic functional relationship between *f_V_*and *ƛ,* and there is a region in which increasing environmental stability (*ƛ*) increases the frequency of generalist genotypes, *f_V_*. How do we explain this?

**Figure 4.**
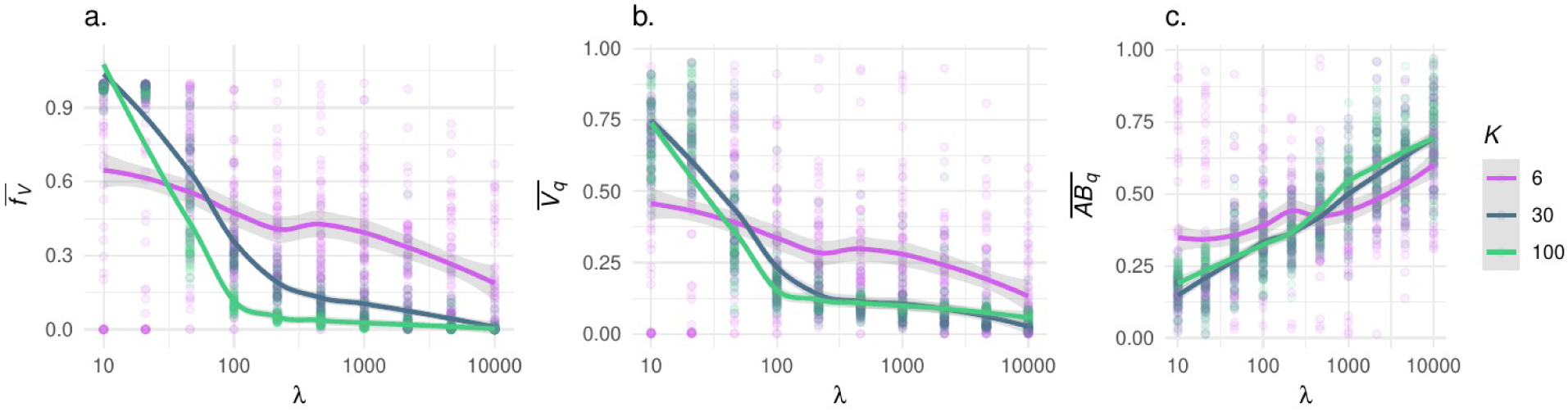
How, when the phenotype space is large, P=100, the number of generations between environmental fluctuations, ƛ, affects (**a**) the mean frequency of generalist genotypes f_V_, (**b**) the mean mutational robustness of genotypes expressing the generalist phenotype V_q_ and (**c**) the mean mutational robustness of specialists, AB_q_. Each point shows the result for one simulation. Lines show loess regressions. Point and line colors denote different values for phenotypic neighborhood size, K. Interpretation: When P is large and K is small, it is unlikely that populations will evolve well-adapted specialists unless the environment is relatively stable. Counterintuitively, increasing environment stability can increase the frequency of mutationally sensitive generalists.

It seems there is an interaction between environmental and mutational robustness. Mean population mutational robustness, *q_i_*, changes little with *ƛ*. But this masks an underlying divergence in *q_i_*between specialist and generalist genotypes (Fig 4b,c.). Specifically, as *ƛ* increases from 10 to 10,000, in generalists *q_i_* drops from 0.45 to 0.13, while in specialists it increases from 0.34 to 0.60. So, when populations with small phenotypic neighborhoods are challenged to adapt to a world in which there are many phenotypic possibilities, environmental stability selects for evolvability in generalists and mutational robustness in specialists. At rates of environmental change that most favor evolvability (100 < *ƛ* < 1000), generalists tends to be more evolvable. And so inside this range of values for *ƛ*, we see an increase in the mean frequency of generalist phenotypes.

Why are generalists more evolvable in such environments? One hypothesis would be that generalists do not persist long enough for selection for mutational robustness to be effective. But closer analysis reveals evidence to the contrary. In fact, over the *ƛ* values at which *f_V_*increases, generalists tend to persist for several thousand generations (Fig. S2.c). Moreover, they tend to evolve lower mutational robustness than their specialist ancestors, and this dynamic becomes more pronounced as *ƛ* increases. If we divide the evolutionary history of a population into phases of generalism (*f_V_* > 0.9) and specialism (*f_V_* <= 0.9), and then calculate the mean change in *q_i_* over the course of a generalist phase, *Δ_qV_*, the mean of this quantity decreases from -0.04 when *ƛ* = 10, to -0.18 when *ƛ* = 10,000 (Fig. S2.a). So, the high evolvability of generalists is not because generalists are ephemeral.

A alternative hypothesis is that selection against mutational sensitivity is less efficient in generalists, since the expected decrease in fitness from non-neutral mutation is less severe. Generalists tend to evolve evolvability via genetic drift. Sure enough, whereas increasing environmental stability, *ƛ,* causes the mean of *Δ_qV_* to become more negative, the standard deviation of *Δ_qV_* remains consistently positive, regardless of environmental stability (Fig. S2.b). And this helps us better understand the tendency for *Δ_qV_* to be negative: Being a generalist does not increase evolvability. The evolution of increased evolvability presages the end of generalism.

So in broad strokes, the inferences of Meyers et al. (26) are robust to relaxations of the their assumptions about genetic architecture. Even if we let mutational sensitivity and phenotypic neighborhood be freely-evolving genotype properties, over much of the explored parameter space, as environments become increasingly stable, we find the same succession of mutationally robust generalists, evolvable specialists, and mutationally robust specialists. On the other hand, we find some new patterns when phenotypic neighborhood sizes are a small fraction of the total number of possible phenotypes. In particular, there can be a negative feedback between mutational sensitivity (1-*q_i_*) and phenotypic neighborhood diversity (*D*) such that more mutationally sensitive genotypes might not be more evolvable. And counter-intuitively, certain increases in environmental stability can cause an increase in the frequency of generalist phenotypes, provided that they are more mutationally sensitive than specialists.

### 4.b. Evolvable cyclical networks of genotypes varying in robustness

#### 4.b.i. Model description

Next, let us extend the Meyers et al. (26) model so that the mutational network is a freely evolvable individual-level property. We mostly follow the original and use the same set of possible phenotypes, **P** = {*A*, *B*, *V*}, the same fitness functions, and the same regimes of fluctuating selection (1 < *ƛ* < 10,000). But we change the mutational process to allow for two kinds of effects. With probability *ρ=*0.5, a mutation transforms the genotype of an individual to one of the adjacent genotypes in their mutational network. With probability 1-*ρ,* a mutation randomly re-assigns the phenotypic mapping of a randomly selected vertex in the individual’s mutational network. To clarify, in the divergent mutational network model of Draghi et al. (25), the network is a population-level property; an individual’s position in that network is determined by their ancestry and the elements of their phenotypic 1-neighborhood. By contrast, with this extension of the Meyers et al. (26) cyclical network model, the network is essentially a higher-order, phenotypic 4-neighborhood, governing what phenotypes are accessible to a genotype via any number of mutations.

We analyze the dynamics of this model with individual-based simulations, with 20K generations for each replicate, and 50 replicates per value of *ƛ*. Model codes are provided in Supplementary Document S4. In each generation, we record the following model statistics: (1) *w*, the mean population fitness, (2) *GP_max_*, the most common genotype-phenotype map, (3) *S^-1^*, the population mean of the inverse richness of mutationally-adjacent phenotypes, which is a measure of mutational robustness, (4) *f_V_*, the mean frequency of individuals expressing a generalist phenotype, and (5) *f_Vk_*, the mean frequency of phenotypic 1-neighborhoods containing the generalist phenotype *p_V_*.

#### 4.b.ii. Model dynamics

In broad strokes, with this model, we find the same *ƛ*-dependent succession of environmentally-robust generalists, evolvable specialists, and mutationally robust specialists (Fig. 5). But we also get a couple of new insights, and a surprise. The first new insight is that when environments turn over rapidly (*ƛ*=10), not only does the generalist phenotype *p_V_* prevail, but mutational networks are enriched for *p_V_* (Fig6a.). In fact, the most common network, occurring at an average frequency of ∼0.43, is composed exclusively of *p_V_* (Fig 6b.). Populations evolve to be both environmentally and mutationally robust. And the mutational robustness observed in this case is more pronounced than anything we see in mutationally-robust specialists in stable environments. The second new insight is that in stable environments (*ƛ*=10,000), mutational robustness is asymmetrical across specialist phenotypes, with higher average robustness for the phenotype that was optimal in the first environmental epoch, *p_A_*. Historical contingency can have long-lasting effects on the evolution of the genotype-phenotype map.

**Figure 5.**
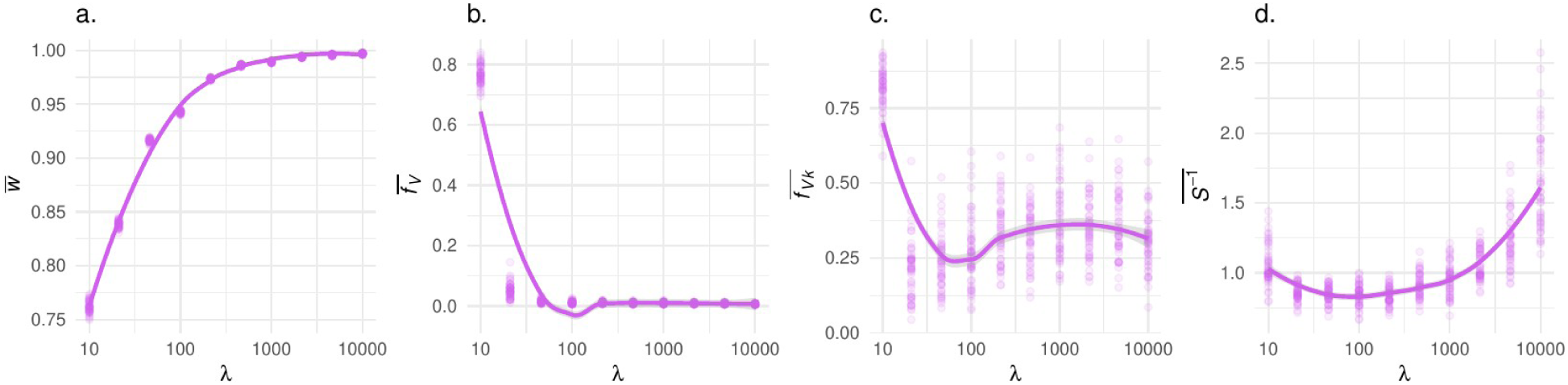
Dynamics of evolving cyclical mutation networks. How the number of generations between environmental fluctuations, ƛ, affects (**a**) the population mean fitness, w-bar, (**b**) the mean frequency of a generalist genotypes f_V_-bar, (**c**) the mean frequency of mutational networks containing the generalist phenotype f_Vk_-bar, and (**d**) the mean phenotypic robustness against mutation, S^-1^. Each point shows the result for one simulation. Lines show loess regressions. The mutation rate μ = 1e-2, and there is an equal probability that a mutation causes an individual to change to an adjacent genotype, or causes a change in the phenotype-assignment of one of the genotypes in the network.

**Fig 6.**
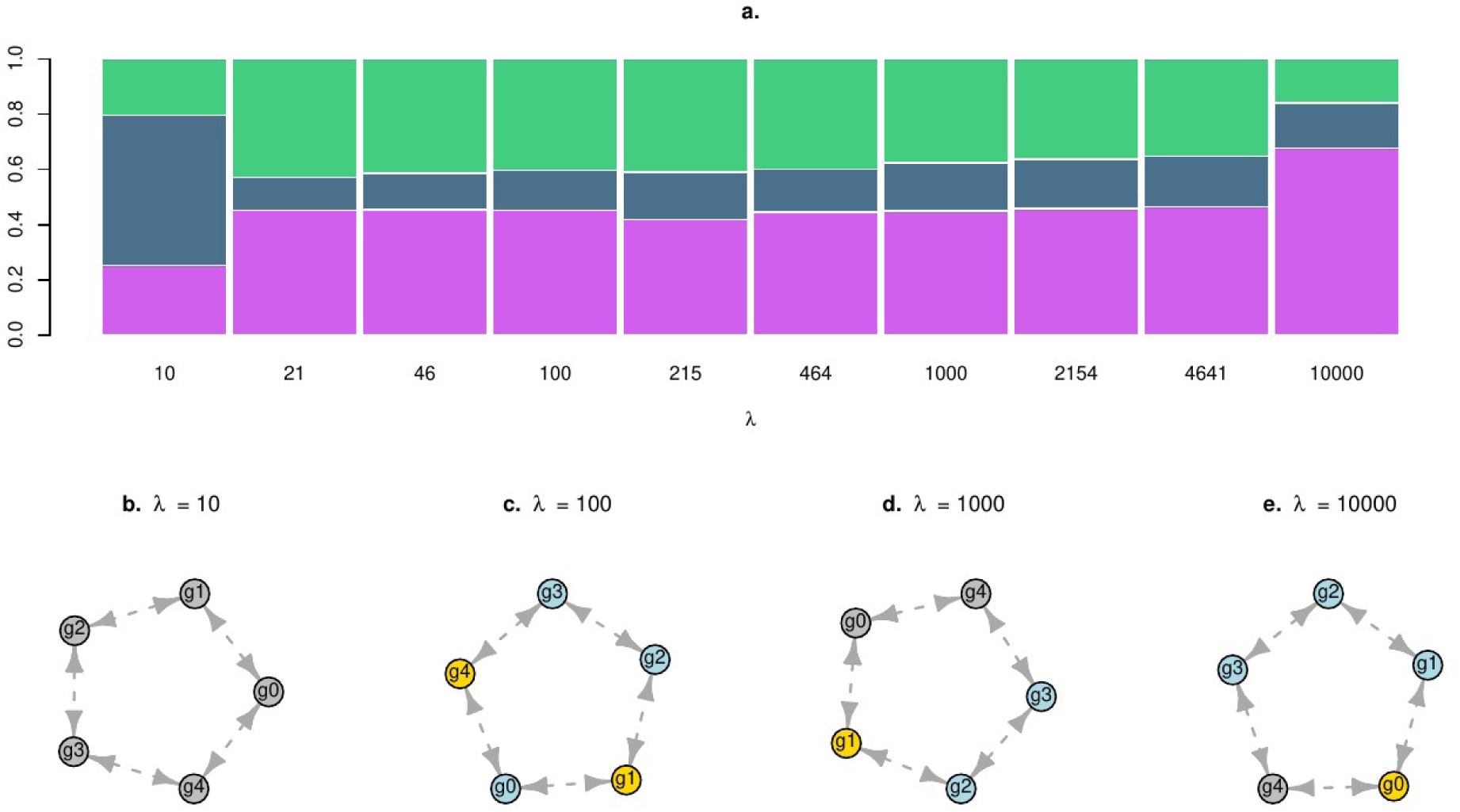
Evolved mutational networks. (**a**) How the number of generations between environmental fluctuations, ƛ, affects the average frequency of each phenotype in the population of mutational networks, with the frequency of p_A_ shown in purple, p_B_ in green, and p_V_ in blue. Below, (**b-e**), the most common mutational network is shown for a subset of ƛ values. Here genotypes mapping to phenotype p_A_ are in light blue, p_B_ in gold, and p_V_ in gray. Interpretation: *When we let cyclical mutational networks evolve in volatile environments, populations evolve to be both mutationally and environmentally robust. Increases in evolvability are realized by evolving networks that are an even mix of specialist phenotypes. And in stable environments, historical contingency can cause long-lasting biases in phenotypic robustness*.

The surprise is that – similar to what we saw for *f_V_*with our extensions of Draghi et al. (25) model – there is a kink in the functional mapping of variation in *ƛ* to *f_Vk_*’, that is, the mean population frequency of genotypes containing the generalist phenotype *p_V_* in their phenotypic 1-neighborhood. A dip in *f_Vk_*’ happens at *ƛ* ∼ 100, where the balance between selection for evolvability and robustness tips most in favor on evolvability; at *ƛ* = 100, in evolved mutational networks, a mapping to *p_V_* is rare, and there is relatively even mix of mappings to *p_A_* and *p_B_* (Fig 7a,c.). Beyond *ƛ* = 100, selection against *p_V_* mappings weakens, as mutational robustness again becomes more valuable. It seems that selection for evolvable specialists more efficiently reduces the likelihood of generalism-causing mutations than does selection for robust specialists.

**Figure 7.**
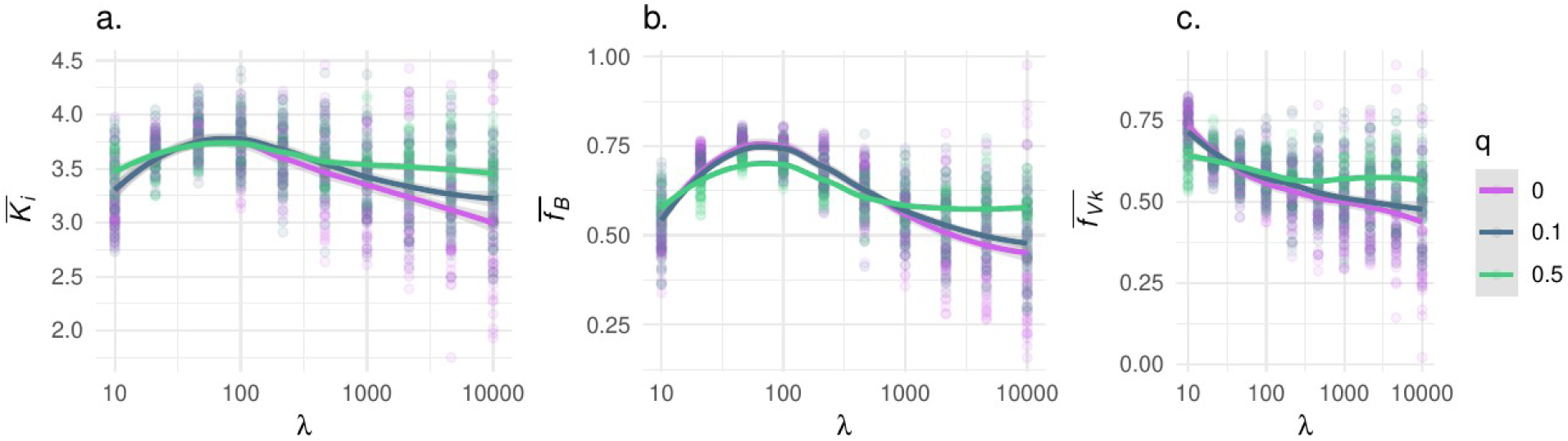
Summary of models where q is fixed and K_i_ is freely evolvable. Panels show how changing the number of generations between environmental fluctuations, ƛ, affects (**a**) mean phenotypic neighborhood size, K_i_-bar; (**b**) mean frequency of phenotypic neighborhoods containing the optimal phenotype for the out-of-phase environmental state, f_B_-bar; and (**c**) mean frequency of phenotypic neighborhoods containing the generalist phenotype, f_Vk-_bar. Each point shows the result for one simulation. Lines show loess regressions. Point and line colors denote different values for q. Interpretation: At intermediate rates of environmental change, populations evolve to maximize evolvability by enriching their phenotypic neighborhoods with phenotypes that are optimal in the out-of-phase environmental state.

## 5. Robustness via phenotypic neighborhood evolution

In the previous sections, we saw that the evolution of mutational robustness, *q_i_*, and how it affects evolvability depends on **k_i_**, the phenotypic neighborhood. In fact, selection on evolvability could affect *q_i_* or **k_i_**. We show this with another model variant (Supplementary Document S6). The model set-up is as described in Section 4.a, for *P*=6, except for three changes: (1) We fix the value of *q* ∈{0.0, 0.1, 0.5}. (2) We let *K* evolve freely; hence, each genotype determines a phenotypic neighborhood size *K_i_*. (3) We change the way that **k_i_** evolves. Specifically, at the start of each simulation, the population is monomorphic for a three-dimensional **k_i_**, with elements randomly sampled from **P**. Then, as the life cycle turns over, mutation can with equal probability add or subtract one element from **k_i_**, with the restriction that 0 < *K_i_* ≤ *P*. We do not prevent redundancy of the elements of **k_i_**. This is to maximize flexibility for the design by selection of an optimal phenotypic neighborhood. To further resolve the model dynamics we also keep track of two additional model statistics, to wit, *f_B_* is the mean frequency of phenotypic neighborhoods that contain the optimal phenotype for the out-of-phase environmental state, and *f_Vk_* is the mean frequency of phenotypic 1-neighborhoods with the generalist phenotype.

With simulation analysis, we find that just as when we fix *K* and let *q_i_* vary across genotypes, populations appear to evolve maximum evolvability at intermediate values for *ƛ* (Fig 7). At intermediate values for *ƛ,* not only is the mean of *K_i_* highest, but so is *f_B_*, the probability that **k_i_** contains the optimal phenotype of the out-of-phase environmental state, and in fact, the latter effect is more pronounced. Averaging across values of *q*, as *ƛ* increases from 10 to 100, there is a 13% increase in *K_i_*, and a concomitant 35% increase in *f_B_*; evolvable populations stack their phenotypic neighborhoods with good options. With increasing *ƛ* we also see a decline in the frequency of the generalist genotype along with *f_Vk_*, that is, the frequency of genotypes that have the generalist phenotype in their neighborhood. These relations are robust to variation in mutational robustness *q*, although when *q* is closer to zero, we see more pronounced reductions of *K_i_*and *f_B_* at extreme values of *ƛ*.

So, there is more than one way to evolve robustness and evolvability. Selection can shape the probability that a mutation will be neutral. Or it can shape the probabilities of the specific phenotypic effects of non-neutral mutation, as has been demonstrated previously with analyses of more complex genotype-phenotype maps (21–23). In fact, the latter would appear to be favored by selection, since tailoring the phenotypic neighborhood can increase the odds of beneficial mutations without also increasing the odds of deleterious ones. Sure enough, if we simulate the evolution of population in which mutational robustness *q_i_*, phenotypic neighborhood size *K_i_*, and phenotypic neighborhood components **k_i_** are all free to vary, then environmental stability, *ƛ,* has little effect on the means of *q_i_* (which tend to be ∼ 0.4) and large effects on means of *K_i_* and *f_B_*, that is, the frequency of phenotypic neighborhood containing the phenotype that is optimal in the out-of-phase environment (Fig. S5). Something to stress here is that for selection on phenotypic neighborhood components to be effective, a genotype-phenotype map needs to have a memory of past selective environments. We made that possible, in an elementary fashion, by implementing a piece-wise mutational routine for adding and removing components of the phenotypic neighborhood, **k_i_**. But more complex genotype-phenotype maps can have better memories (22,24).

## 6. Discussion

Here we addressed this question: How does the robustness of a discrete-state phenotype relate to evolvability when it can vary across genotypes and phenotypes, against variation in both the genetic and external environment? We improved our intuition by considering some simple models in which robustness is expressed as *q*, the probability that some perturbation does not affect the phenotype, and if a perturbation does affect the phenotype, such effects are mediated by **k**, the neighborhood of accessible phenotypes. For ease of discussion, let us call these *qk* (pronounced “cuke”) models. By merging and extending previously-developed *qk* models (25,26), we saw that their inferences are robust and their scope of applicability is broader than has been appreciated; both tell us as much about the evolution of environmental robustness as mutational robustness. We also gained two additional key insights. First, counter-intuitively, in certain genetic and environmental contexts, reducing the rate of environmental change can lead to selection for more environmentally robust genotypes, that is, generalists. This is because generalists tend to evolve lower levels of mutational robustness. Second, although the analysis of *qk* models has focused on *q*, the probability of non-neutral mutation, selection for increased evolvability works preferentially on *k*, the neighborhood of mutationally-accessible phenotypes. This is because evolability via phenotypic neighborhood evolution comes without an increase in mutational load. That selection can affect the distribution of mutational effects has been shown before with analysis of more complex genotype-phenotype models (19,24,33,34). But to my knowledge, this is the clearest demonstration that increasing evolvability by evolving the phenotypic neighborhood evolution is a better option than evolving increased mutational sensitivity.

Another new insight is that the interaction between different kinds of robustness can cause counter-intuitive evolutionary dynamics. Specifically, in complex environments, increasing environmental stability can cause an increase in the frequency of environmentally-robust genotypes, that is, generalists. We saw that this corresponds to a divergence in the mutational sensitivities of generalists and specialists, and we traced this divergence to a neutral evolutionary process; the evolution of evolvability in generalists is accidental, but predictably so. This is yet another example of how robustness can increase evolvability by facilitating the accumulation of cryptic genetic diversity (32,35).

There are many things that *qk* models of the evolution of robustness and evolvability can not do. In particular, robustness is just one of several properties of a genotype-phenotype map that can affect evolvability, and even the effects of robustness can depend on other facets of genetic architecture. Therefore, a richer understanding of the evolution of evolvability requires more complex models genotype-phenotype maps. Other germane properties include a developmental system’s criticality, that is, the robustness of its robustness (19,36), as well as its capacity for plasticity and cryptically-neutral genetic diversity (18,20,22), and its capacity to adaptively remember and generalize from past selective environments (23,24,37). More complex genotype-phenotype map models can also yield insights into the specific design features that determine map properties such as robustness, evolvability and memory. A burgeoning body of research has already revealed several such network features (38), such as modularity (33,39), excitation (17), and scale-free degree distribution (40,41). Researchers have also begun to detect important network motifs affecting robustness and evolvability, such as freed-back and feed-forward loops (42). So, the analysis of more complex genotype-phenotype map models promises to yield a more multi-faceted and mechanistic understanding of the evolution of evolvability. Nevertheless, by ignoring the genetic architectural details that might make one genotype-phenotype map more robust than another, *qk*-models throw into sharp relief the effects of robustness *per se* on evolvability.

## Acknowledgments

Thanks to Jeremy Draghi, and Ben Normark for providing helpful comments on a draft of the manuscript.

