## Supplementary Figures for "A robust relationship between robustness and evolvability"

1 Genetic Robustness and Evolvability

### Supplementary Figures

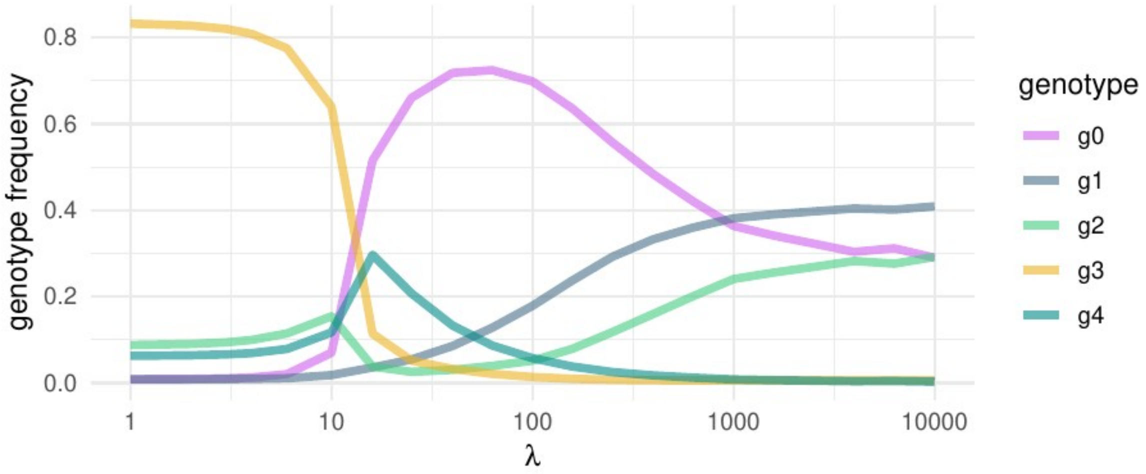


**Figure S1.** *Reproduction of the analysis of the Meyers et al. (26) with R code provided as Supplementary File S1. Environmental stability, λ̃, is given on a log scale on the x-axis. Curves show the mean frequency of each genotype when the environment is in state a.* Interpretation*: In volatile environments, generalists predominate. In stable environments mutationally robust genotypes rule. In environments that change at intermediate frequency, the population evolves to maximized evolvability; it becomes an evolvability specialist.*

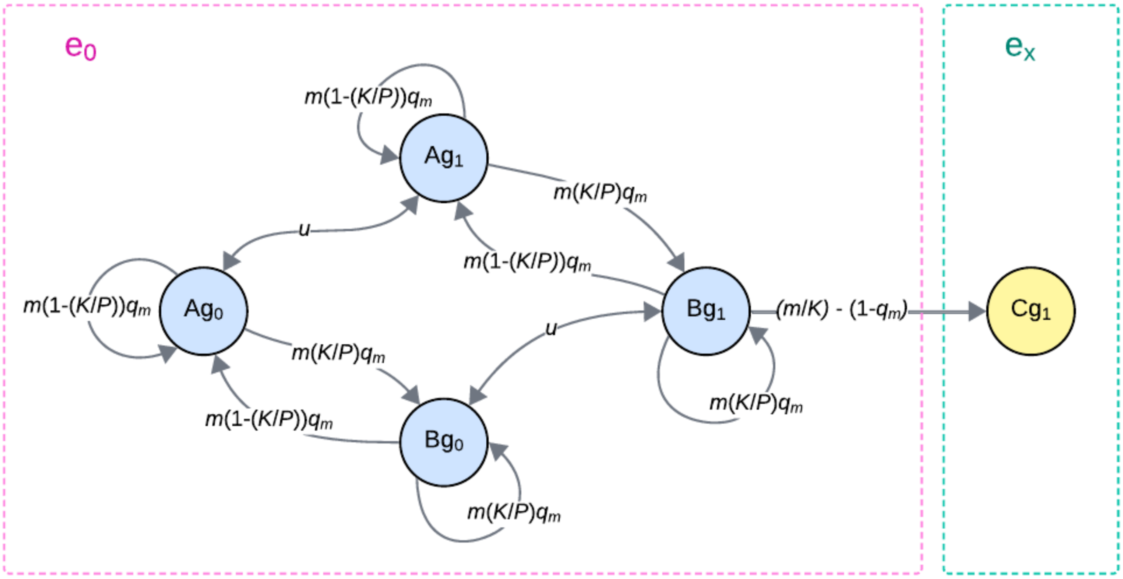


**Figure S2.** *Flow diagram of model relating environmental robustness to evolvability. Vertices represent model variables, and edges show how one variable affects another, or if an edge exits and enters the same vertex, how a variable affects itself. Extending the scheme of Draghi et al. (25), we divide the population of individuals into compartments, based on whether they, after $t_x$, (C) occupy and have high fitness in environment $e_x$, or (B) have $e_x$ in their environmental neighborhood – a component of the decomposed phenotypic neighborhood* $\mathbf{k}_i$*, which combines environmental and genetic neighborhoods – or (A) do not have $e_x$ in their environmental neighborhood, with compartments A and B further divided by genotype. So, each model variable is the number (or proportion) of a particular genotype in a particular environment type. The expression on each edge gives the rate at which individuals flow from one compartment to another. Take for an example the edge flowing from variable $B_{g1}$ to $C_{g1}$. If an individual is in a B compartment, that means they have environment $e_x$ in their environmental neighborhood. So the probability that an individual in a B compartment with the $g_1$ genotype migrates to environment $e_x$ is equal to the migration rate m, times the probability that a non-neutral a migration lands them environment $e_x$, 1/K, minus the probability that a migration is not neutral ($1$-$q_m$).*

3 Genetic Robustness and Evolvability

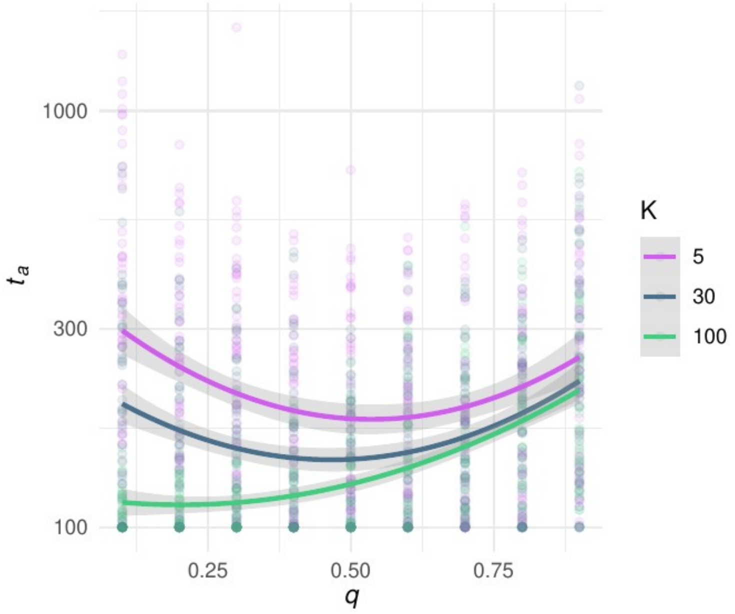


**Figure S3.** *The effect of migrational robustness, $q_m$, on evolvability, $t_a$, depends on the difference between the number of possible phenotypes, K, and the size of the phenotypic neighborhood P. Curves are loess regressions; shaded areas give 95% confidence intervals.* Interpretation*: Unless all phenotypes are accessible by one migrational step, evolvability is maximized at intermediate levels of migrational robustness.*

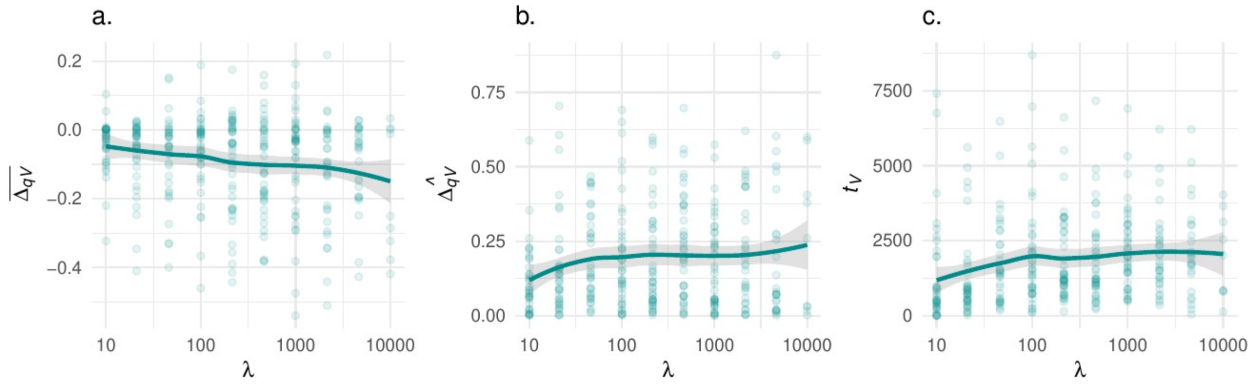


**Figure S4.** *The evolution of evolvable generalists. How the number of generations between environmental fluctuations,* $\bar{\lambda}$*, affects (**a**) the mean change in mutational robustness over the course of an evolutionary phase of generalism,* $\Delta_{qV}$*, (**b**) the standard deviation of that same quantity, and (**c**) the mean duration of an evolutionary phase of generalism. Each point shows the value for one simulation. Lines show loess regressions.* Interpretation: *Generalists tend to persist for thousands of generations, and to evolve lower mutational robustness than their specialist ancestors. But the latter effect is likely due to neutral genetic processes.*

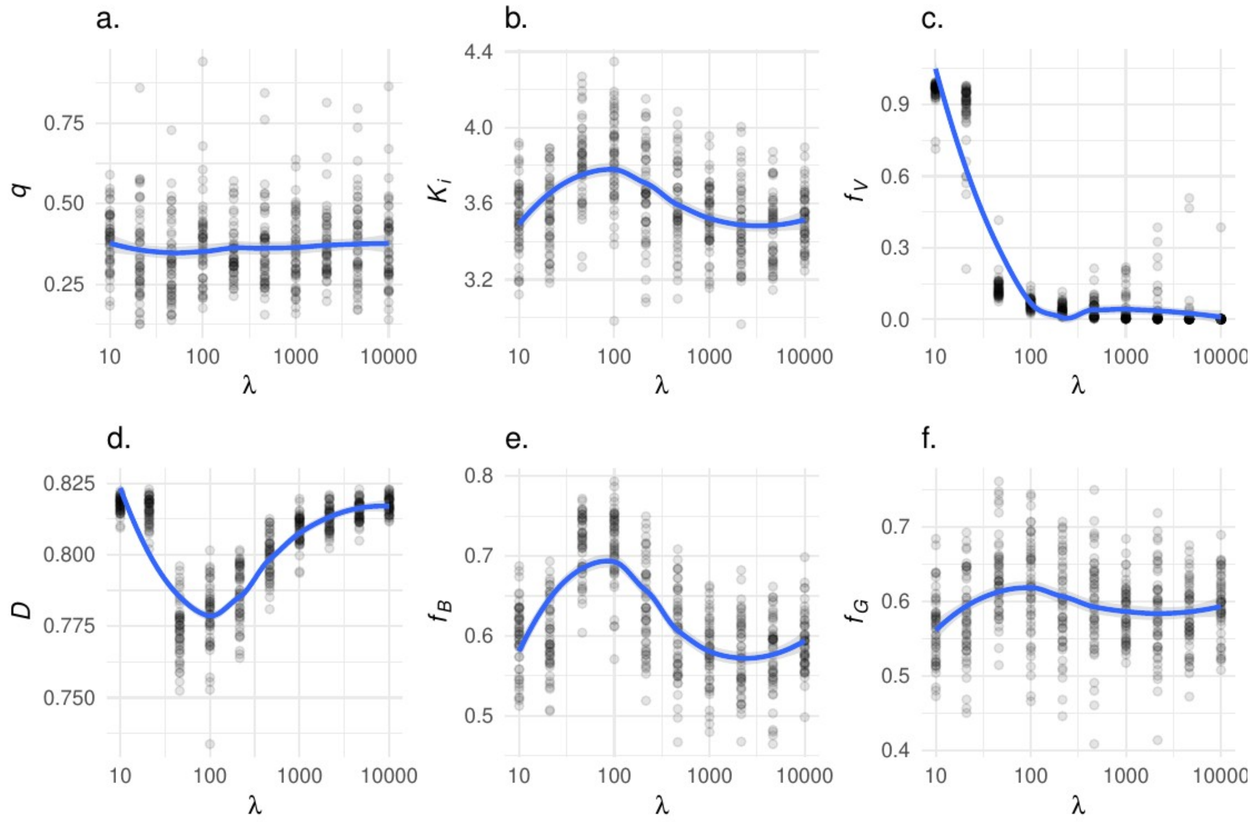


**Figure S5.** *Summary of models where qi and* $K_i$ *are both freely evolvable. How changing the number of generations between environmental fluctuations,* $\tilde{\lambda}$, *affects (**a**) mean mutational robustness* $q_i$*-bar, (**b**) mean phenotypic neighborhood size,* $K_i$*-bar; (**c**) mean frequency of generalist genotypes* $f_V$*-bar, (**d**) mean population diversity of phenotypic neighborhoods, D-bar; (**e**) mean frequency of phenotypic neighborhoods containing the optimal phenotype for the out-of-phase environmental state,* $f_B$*-bar; and (**f**) mean frequency of phenotypic neighborhoods containing the generalist phenotype,* $f_{Vk}$*-bar. Each point shows the result for one simulation. Lines show loess regressions. Point and line colors denote different values for q.* Interpretation*: Selection for evolvability favors changes in phenotypic neighborhood components over mutational robustness.*
